# Associations between oral microbiota pathogens and elevated depressive and anxiety symptoms in men

**DOI:** 10.1101/2024.08.10.606819

**Authors:** Fannie Kerff, Julie A Pasco, Lana J Williams, Felice N Jacka, Amy Loughman, Samantha L Dawson

## Abstract

**Background:** Systemic inflammation has been associated with depression. Certain oral bacterial species contribute to extra oral inflammation, however their potential association with mental disorders remains unclear. This study investigated the associations between oral microbiota pathogens and depressive and anxiety symptoms.

**Methods:** Data came from 436 men from the Geelong Osteoporosis Study. Using 16S rRNA gene sequencing, we characterized the oral microbiota composition and created a composite variable representing the prevalence of known oral pathogens, *Porphyromonas gingivalis*, *Treponema denticola*, *Fusobacterium nucleatum*, and *Prevotella nigrescens*. Binary variables were created representing elevated depressive and anxiety symptoms using the Hospital Anxiety and Depression Scale. Associations between oral pathogens and elevated depressive/anxiety symptoms were analyzed using logistic regression adjusted for confounders (age, socio-economic status, diet, smoking, alcohol, exercise, obesity and hypertension).

**Results:** We found limited evidence for an association between the pathogen composite and elevated depressive (N = 39) but not anxiety (N = 66) symptoms. Moreover, some of the comprising species, *P. nigrescens* (OR 1.61 (95% CI 1.21, 2.13)) and *F. nucleatum (subsp. animalis ATCC 51191* (1.54 (1.13, 2.11)), *subsp. animalis* (1.47 (1.05, 2.08)) and *subsp. vincentii* (1.47 (1.06, 2.04))) were associated with elevated depressive symptoms. Exploratory analyses revealed several other taxa associated with depression and anxiety.

**Limitations:** This study focuses on mental health symptoms and not diagnoses in men, and the results may not be generalizable to women.

**Conclusions:** We identified several taxa within the oral microbiota that were associated with depressive and anxiety symptoms. Further replication and mechanistic studies are needed.

**Highlights:**

- A composite of oral pathogens is associated with depressive symptoms in adult men
- Associations between *P. nigrescen*s and other species and elevated depressive symptoms
- Associations remain following adjustment for confounders
- Oral microbial diversity was not associated with depressive or anxiety symptoms
- Fewer associations between oral microbial composition and anxiety than depression

## 1. Introduction

Major depressive disorder and anxiety disorders are common mental disorders that constitute a significant global burden, representing the 13^th^ and 24^th^ leading causes of global disability-adjusted life years, respectively (GBD 2019 Mental Disorders Collaborators, 2022). Cross-sectional studies associate systemic inflammation with elevated depressive and anxiety symptoms (Simpson et al., 2020). Additionally, longitudinal studies indicate that higher levels of inflammation (IL-6, C-reactive protein) precede the onset of depressive symptoms, depressive disorders, psychotic experiences and psychotic disorder (Bell et al., 2017; Khandaker et al., 2014; Lamers et al., 2019; Pasco et al., 2010). The gut microbiota has emerged as a potential mediator in common mental disorders, interacting via the microbiota-gut-brain axis with stress and inflammatory pathways (Marx et al., 2021). The oral microbiota is directly connected to the gut microbiota via the gastrointestinal tract (Park et al., 2021) and may have a direct route from the oral cavity to the brain, via the trigeminal nerve (Riviere et al., 2002) and/or the blood-brain barrier (Liu et al., 2023). The association between the oral microbiota and common mental disorders needs further investigation.

A small number of studies have investigated differences in the oral microbiota in those with depression compared to controls (Kohn et al., 2020; Simpson et al., 2020; Wingfield et al., 2021; Yolken et al., 2021). These typically have small sample sizes, and not all studies report differences (Yolken et al., 2021). Healthy non-smoking adults with higher distress exhibited greater salivary microbiota diversity compared to those with lower distress (Kohn et al., 2020). Whereas in another study in adolescents, specific bacterial taxa in the oral microbiota were associated with anxiety and depressive symptoms but there were no differences in diversity (Simpson et al., 2020). The family, order, and phylum of the genus *Treponema*, namely *Spirochaetaceae*, *Spirochaetales*, and *Spirochaetes*, were all positively associated with the severity of depressive and anxiety symptoms in adolescents. Compared to controls matched for age, sex, and -where possible-smoking status, young adults with depression had a significantly greater abundance of *Prevotella nigrescens* and the genus *Neisseria*, as well as differences in beta (but not alpha) diversity (Wingfield et al., 2021).

Alzheimer’s disease shares some similar inflammatory and neuroimmune pathophysiology with common mental disorders (Lau et al., 2023; Liu et al., 2023), with particular oral pathogens implicated in both. For example, *Porphyromonas gingivalis* (Dominy et al., 2019) and their lipopolysaccharides (Poole et al., 2013) infected the cerebrospinal fluid and the brain of post-mortem Alzheimer’s disease patients. A seven-times greater load of *Spirochaetes* was also detected in brains from people with Alzheimer’s disease (Miklossy, 2011), suggesting potential interactions of *P. gingivalis* with pathogenic *Treponema* taxa such as *Treponema denticola* (Kin et al., 2020; Nieminen et al., 2018). Moreover, the presence of the *Treponema* genus in Alzheimer’s patients’ trigeminal nerves suggested one of its potential routes of access to the brain (Riviere et al., 2002). Regarding bipolar affective disorders, depressive phases have been associated with elevated levels of *Aggregatibacter actinomycetemcomitans* and *P. gingivalis* (Cunha et al., 2019). Furthermore, *Klebsiella pneumoniae* and *Fusobacterium nucleatum* have been implicated in inflammation and may be associated with extra oral inflammatory comorbidities (Hajishengallis and Chavakis, 2021).

This study aimed to investigate the association between the oral microbiota, focusing on the candidate species identified above, and elevated depressive and/or anxiety symptoms in participants from an Australian cohort.

## 2. Methods

This study was pre-registered in May 2023 on the Open Science Framework platform (https://osf.io/wcfkp/). The Human Research Ethics Committee at Barwon Health approved the study (00/56).

### 2.1 Study design and sample

The Geelong Osteoporosis Study (GOS) is an ongoing, observational cohort study of adults randomly selected from the electoral roll. Inclusion criterion was a listing as a resident of the Barwon Statistical Division in south-eastern Australia; residency in the region for less than six months and inability to provide informed consent necessitated exclusion (Pasco et al., 2012). This study utilizes cross-sectional data from 436 men who provided an oral sample as part of the 15-year male follow-up, conducted from 2016 to 2020. Women were not included in this analysis, as oral samples were not available at the time of writing. Participants were included in the analysis dataset if they had complete data for the exposure (gum swab) and outcome (depressive and anxiety symptoms). Men were excluded if they had a past-year diagnosis of severe disease (including cancer, heart, and brain diseases), due to the potential non-trivial effects of these conditions on the oral microbiota and depressive/anxiety symptoms.

### 2.2 Oral microbiota exposure measures

#### Collection and processing

Oral microbiota samples were collected via swabs of the upper and lower gum-line using sterile cotton tips. The samples were stored in a −80°C freezer located at the University Hospital Geelong, Barwon Health. We used bacterial 16S ribosomal RNA (16S rRNA) gene-based next generation sequencing (NGS) to profile the bacterial composition. Library preparation and sequencing was conducted at Charles River Laboratory (Australia), using two sets of 16S primers 27F-336R (V1-V2) and 341F-785R (V3-V4). For sequencing, the NovaSeq SP 500 cycle flow cell (NV058 and NV058RE2) 251 | 10 | 10 | 251 was used.

Oral microbiota data were analyzed using the R package *phyloseq* (v1.44.0). No filtering of low abundant or low prevalent features was performed for the main analysis, given the need to extract the maximum number of Operational Taxonomic Units (OTUs) from candidate pathogens in the samples. After agglomerating the oral taxa at the species level, a Centered Log Ratio (CLR) transformation was performed on all oral taxa as per best practice (Gloor et al., 2017). The CLR transformation of the data was performed using the CLR transform function of the *Tjazi* package (Bastiaanssen et al., 2023), reflecting how OTUs perform relative to the per-sample average (Quinn et al., 2019). The zero imputation was implemented using the Martın-Fernandez et al. method, meaning replacing all zeros with 65% of the detection limit, threshold minimizing the distortion in the covariance structure (Lubbe et al., 2021).

#### Oral pathogen composite

To investigate candidate oral species hypothesized to be associated with elevated depressive and/or anxiety symptoms, a multiple-species composite score was computed following a compositional data analysis approach, henceforth referred to as pathogen composite. The rationale behind employing this method lies in the understanding that microbes operate in functional guilds rather than in isolation (Wu et al., 2021), and microbiota studies are often underpowered (Kers and Saccenti, 2022). All components of the composite are interdependent features that cannot be fully understood in isolation (Quinn et al., 2019).

To build this pathogen composite, the CLR transformed abundances of the OTUs corresponding to candidate oral species were retrieved and summed into one value per sample, reflecting its pathogen composite score. The candidate oral pathogens listed in the pre-registration of this study included *P. gingivalis*, *F. nucleatum*, *T. denticola*, *A. actinomycetemcomitans*, and *K. pneumoniae*. Since then, evidence of an association between oral *P. nigrescens* and clinical depression has arisen (Wingfield et al., 2021), and therefore these were also included in the pathogen composite.

#### Microbial diversity

The richness, diversity and degree of variation of the oral microbiota were quantified from the raw count table using the Alpha diversity indices Chao1, Simpson, and Shannon entropy. Following a prevalence filtering set at 1% to remove rare taxa, beta diversity was evaluated using principal component analysis based on the Aitchison distance (Aitchison, 1982) (i.e., Euclidean distance over CLR-transformed values (Bastiaanssen et al., 2023)).

### 2.3 Outcome measures

#### Depressive and anxiety symptoms

Depressive and anxiety symptoms were assessed using the 14-item Hospital Anxiety and Depression Scale (HADS) questionnaire, a self-report measure of symptom frequency over the past week (Bjelland et al., 2002). HADS scores 0–7 represent “normal”, 8–10 “mild”, 11–14 “moderate”, and 15–21 “severe” symptoms (Pais-Ribeiro et al., 2018). The cut-off score at ≥ 8 for both HADS-A and HADS-D has been shown to provide a balance between sensitivity and specificity (i.e., sensitivities and specificities for both subscales are approximately 0.8) and provides a suitable screening threshold (Bjelland et al., 2002).

#### Primary outcome

We created a binary case group defined as HADS-D score of ≥ 8 representing “elevated depressive symptoms”, in line with consensus in previous literature (Bjelland et al., 2002); scores less than 8 represented “minimal depressive symptoms”.

#### Secondary outcome

Similarly, HADS-A scores were dichotomized, with HADS-A scores ≥ 8 representing “elevated anxiety symptoms” (Bjelland et al., 2002); scores under 8 represented “minimal anxiety symptoms”.

### 2.4 Covariates

The associations between the oral microbiota and the presence of elevated depressive/anxiety symptoms were modelled using a causal inference modelling framework for observational data (Hernán et al., 2022). A directed acyclic graph was used to identify potential confounders for common mental disorders (Otte et al., 2016) and the oral microbiota: age (Burcham et al., 2020), socio-economic status (SES) (Belstrøm et al., 2014), diet (Liu et al., 2022), smoking (Willis et al., 2022; Wu et al., 2016), current alcohol intake (Fan et al., 2018), physical activity (Uchida et al., 2021), and the presence of a chronic disorder, with most evidence on obesity (Burcham et al., 2020; Wu et al., 2018) and hypertension (Willis et al., 2022) (included in the pre-registration; Supplementary Fig. 1). SES was determined using the deciles (1-10) of the Index of Relative Socio-economic Advantage and Disadvantage (IRSAD), a component of the Socio-Economic Indexes for Areas (Australian Bureau of Statistics). To measure the inflammatory potential of participants’ diets, the Energy-adjusted Dietary Inflammatory Index (E-DII) was computed (Li et al., 2021; Shivappa et al., 2014) based on responses from the Dietary Questionnaire for Epidemiological Studies (DQES) developed by the Cancer Council Victoria (Giles and Ireland, 1996). A lifestyle risk score (0-5) was defined using the information from smoking, current alcohol intake and physical activity (Xu et al., 2021); a detailed description of its computation is included in Supplementary Table 1. Obesity was defined as having a body mass index of ≥ 30 (World Health Organization); hypertension was self-reported by participants. Missing covariate data were imputed to the median of the cohort.

### 2.5 Statistical analysis

Statistical analyses were performed using R (v4.3.0).

#### Main analysis

Logistic regressions (R package *stats* (v4.3.0)) were computed to measure the association between the oral pathogens, first together as a composite, then individually, and elevated depressive and anxiety symptoms. Both unadjusted and adjusted models were created, the latter controlling for covariates (age, SES, diet, lifestyle risk (smoking, alcohol intake, and exercise), obesity, and hypertension). The logistic regression models measured how a change of one standard deviation in the oral bacteria exposure (CLR transformed) was related to the odds of having elevated depressive or anxiety symptoms.

#### Exploratory analysis

Alpha diversity metrics were calculated using the R package *Tjazi* (0.1.0.0). To investigate beta diversity, an analysis of similarities (ANOSIM) was performed using the R package *vegan* (v 2.6.4).

After applying a prevalence filtering of 10%, differential abundance analyses between all oral microbiota species in the samples and elevated depressive and anxiety symptoms were performed using the *MaAsLin2* package (Mallick et al., 2021), both in unadjusted and adjusted models. *MaAsLin2* parameters included a CLR transformation, to align with the primary analyses, and a statistical significance set at a p-value < 0.05 and a q-value < 0.1, after applying a Benjamini-Hochberg adjustment for multiple testing (Benjamini and Hochberg, 1995).

## 3. Results

### 3.1 Descriptive characteristics of the included participants

Four hundred and thirty-six participants were included in the study (flow chart in Supplementary Fig. 2; details of the excluded conditions in Supplementary Table 2). Participants had a median age of 62 years and a median IRSAD decile score of 6, indicating moderate levels of relative advantage and disadvantage (Table 1). The median E-DII was −0.07, indicating a diet that is neither distinctly anti-inflammatory nor pro-inflammatory, and the average lifestyle risk was 2 out of a possible score of 0-5, representing a relatively low risk sample regarding smoking, alcohol use, and lack of physical activity. Obesity was recorded in 26.2% of participants and 33.7% had hypertension. Thirty-nine men (8.9%) had elevated depressive symptoms, and 66 (15.1%) had elevated anxiety symptoms (Table 1).

**Table 1.** Participant characteristics. Description of eligible participants, and the depressive and anxiety symptoms’ sub-groups. The pathogen composite reflects the combined load of *P. gingivalis*, *T. denticola*, *F. nucleatum subsp. vincentii ATCC 49256*, *F. nucleatum subsp. vincentii 3_1_36A2*, *F. nucleatum subsp. vincentii*, *F. nucleatum subsp. animalis 7_1*, *F. nucleatum subsp. polymorphum*, *F. nucleatum subsp. animalis ATCC 51191*, *F. nucleatum subsp. animalis* and *P. nigrescens ATCC 33563*. Median (IQR); n (%); SES, Socio-Economic Status; IRSAD, Index of Relative Socio-economic Advantage and Disadvantage; E-DII, Energy-adjusted Dietary Inflammatory Index; HADS-D/A, Depression/Anxiety score from the Hospital Anxiety and Depression Scale questionnaire; CLR, Centered Log Ratio; IQR, interquartile range.

|  | All participants<br>(N=436) | Minimal<br>Depressive<br>Symptoms (N=397) | Elevated<br>Depressive<br>Symptoms (N=39) | Minimal<br>Anxiety<br>Symptoms (N=370) | Elevated<br>Anxiety<br>Symptoms (N=66) |
| --- | --- | --- | --- | --- | --- |
| Age (y) | 62 (50, 71) | 62 (51, 71) | 57 (50, 74) | 63 (52, 72) | 54 (44, 68) |
| SES: IRSAD deciles | 6 (4, 7) | 6 (4, 8) | 5 (3, 7) | 6 (4, 7) | 6 (3, 7) |
| Diet: E-DII | -0.07 (-0.90, 0.63) | -0.07 (-0.90, 0.62) | -0.23 (-0.91, 0.68) | -0.11 (-0.98, 0.58) | 0.04 (-0.45, 0.81) |
| Lifestyle risk:<br>smoking, alcohol<br>and exercise | 2 (2, 3) | 2 (2, 3) | 3 (2, 3) | 2 (2, 3) | 3 (2, 3) |
| Chronic disorder:<br>Obesity | 114 (26.15%) | 103 (25.9%) | 11 (28.2%) | 96 (25.9%) | 18 (27.3%) |
| Chronic disorder:<br>Hypertension | 147 (33.72%) | 131 (33.0%) | 16 (41.0%) | 126 (34.1%) | 21 (31.8%) |
| Pathogen composite<br>(CLR) | 9 (2, 17) | 9 (2, 16) | 11 (6, 18) | 9 (2, 17) | 10 (4, 17) |
| Elevated depressive<br>symptoms:<br>HADS-D ≥ 8 | 39 (8.94%) | | | 20 (5.4%) | 19 (28.8%) |
| Elevated anxiety<br>symptoms:<br>HADS-A ≥ 8 | 66 (15.14%) | 47 (11.8%) | 19 (48.7%) | | |

### 3.2 Oral pathogen composite

The swab samples included information on 900 unique OTUs. We included in the composite the candidate oral species that were detected in the samples. *K. pneumoniae* species were not included as no OTU assigned to them were detected; similarly, we did not include *A. actinomycetemcomitans* and *F. nucleatum subsp. fusiforme ATCC 51190* species as these were poorly detected in the samples and not correlated with the composite (details in Supplementary Table 3; correlation matrix in Supplementary Fig. 3). After retrieving the OTUs of the candidate pathogens and computing their correlations, the pathogen composite reflected the loads of *P.gingivalis*, *T. denticola*, *F. nucleatum* (*F. nucleatum subsp. vincentii ATCC 49256*, *F. nucleatum subsp. vincentii 3_1_36A2*, *F. nucleatum subsp. vincentii*, *F. nucleatum subsp. animalis 7_1*, *F. nucleatum subsp. polymorphum*, F*. nucleatum subsp. animalis ATCC 51191* and *F. nucleatum subsp. animalis*) and *P. nigrescens* (*P. nigrescens ATCC 33563*) species. The CLR transformation of the data resulted in a near normal distribution of the pathogen composite score (Supplementary Fig. 4).

### 3.3 Associations between the oral pathogen composite and elevated depressive and anxiety symptoms

There was limited evidence for an association between the pathogen composite score and the presence of elevated depressive symptoms, both in unadjusted and adjusted models (OR (odds ratio) 1.35 (95% CI (95% confidence interval) 0.974, 1.87), Fig. 1). There were no associations observed between the oral pathogen composite score and elevated anxiety symptoms.

**Fig. 1.**
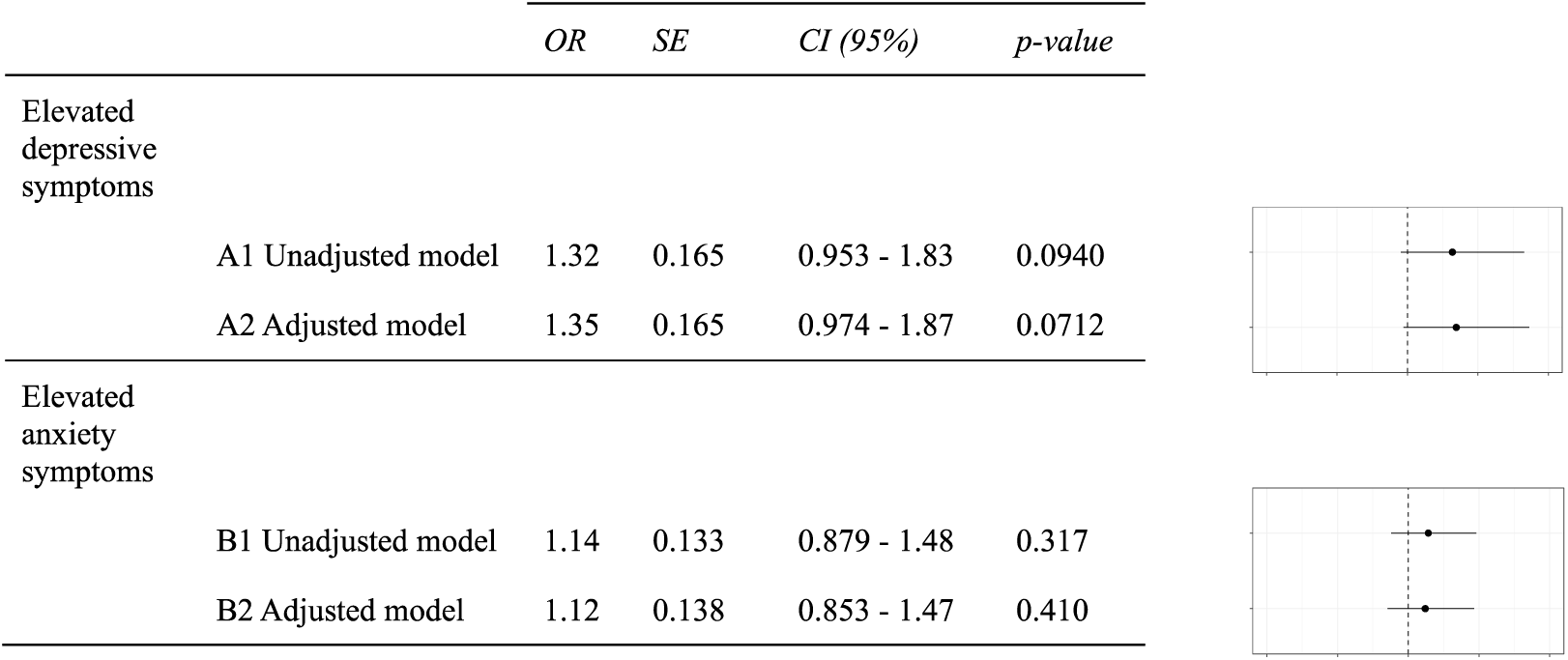
Logistic regressions result for the main analysis: association between the oral pathogen composite and elevated depressive and anxiety symptoms. Limited evidence of associations between the pathogen composite and elevated depressive and anxiety symptoms. Adjusted models control for confounders: age, SES, diet, lifestyle risk, obesity and hypertension. The pathogen composite reflects the combined load of *P. gingivalis*, *T. denticola*, *F. nucleatum subsp. vincentii ATCC 49256*, *F. nucleatum subsp. vincentii 3_1_36A2*, *F. nucleatum subsp. vincentii*, *F. nucleatum subsp. animalis 7_1*, *F. nucleatum subsp. polymorphum*, *F. nucleatum subsp. animalis ATCC 51191* and *F. nucleatum subsp. animalis* and *P. nigrescens ATCC 33563*. OR, Odds Ratio estimate; SE, Standard Error; CI (95%), 95% Confidence Interval; SES, Socio-Economic Status.

### 3.4 Associations between oral species within the pathogen composite and elevated depressive and anxiety symptoms

Several candidate oral pathogens within the composite were positively associated with elevated depressive symptoms (Fig. 2). Species *P. nigrescens ATCC 33563* were significantly more abundant in participants with elevated depressive symptoms, compared to those with minimal depressive symptoms, both in the unadjusted and adjusted models, the latter controlling for age, SES, diet, lifestyle risk, obesity, and hypertension; an increase of one standard deviation of *P. nigrescens* (CLR-transformed) was associated with a 61% increase in the odds of having elevated depressive symptoms (1.61 (1.21, 2.13), p < 0.001, Fig. 2; Supplementary Fig. 5). Three species of *F. nucleatum* were also positively associated with elevated depressive symptoms both in the unadjusted and adjusted models: *F. nucleatum subsp. animalis ATCC 51191* (1.54 (1.13, 2.11), p < 0.01, Fig. 2; Supplementary Fig. 5), *F. nucleatum subsp. Vincentii* (1.47 (1.06, 2.04), p < 0.05, Fig. 2; Supplementary Fig. 5), and *F. nucleatum subsp. Animalis* (1.47 (1.05, 2.08), p < 0.05, Fig. 2; Supplementary Fig. 5). The abundances of *P. gingivalis*, *T. denticola* and the other *F. nucleatum* species were not significantly associated with elevated depressive symptoms (all p > 0.05, Fig. 2). None of the candidate oral species was significantly associated with elevated anxiety symptoms, compared to minimal anxiety symptoms, in unadjusted or adjusted models (all p > 0.05, Supplementary Fig. 6).

**Fig. 2.**
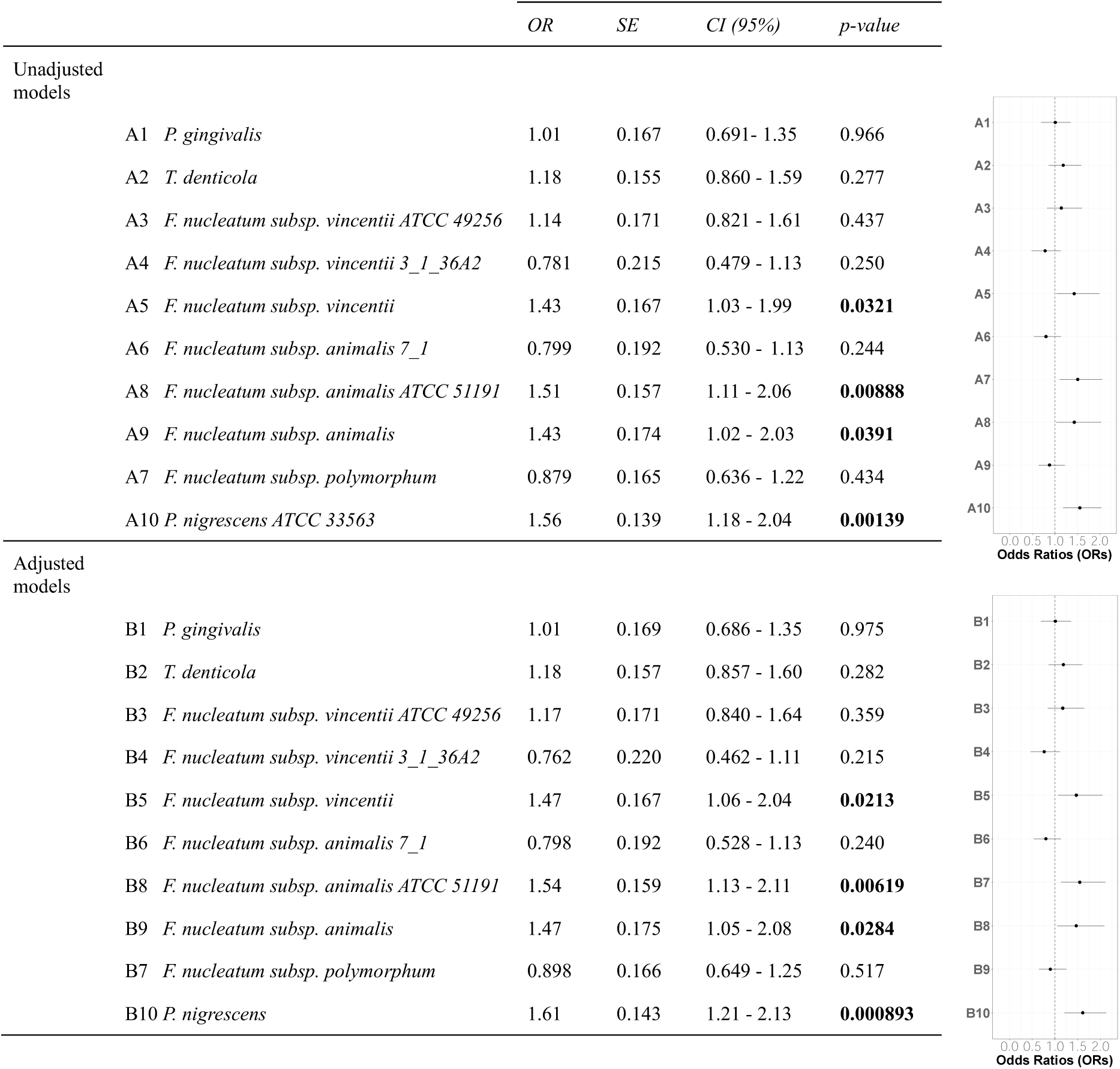
Logistic regressions result for taxa within the composite and elevated depressive symptoms. Significant positive associations between *P. nigrescens*, *F. nucleatum subsp. vincentii*, *F. nucleatum subsp. animalis ATCC 51191*, *F. nucleatum subsp. animalis* and elevated depressive symptoms, both in adjusted and unadjusted models. No evidence of associations with the other species. Adjusted models control for confounders: age, SES, diet, lifestyle risk, obesity and hypertension. OR, Odds Ratio estimate; SE: Standard Error; CI (95%): 95% Confidence Interval; SES, Socio-Economic Status.

### 3.5 Association between oral microbiota diversity and elevated depressive and anxiety symptoms

There were no differences in alpha diversity (Chao1 index, Shannon entropy, or Simpson index) between participants with elevated and minimal depressive symptoms (all p > 0.05, Fig. 3**A-C**), nor anxiety symptoms (all p > 0.05, Supplementary Fig. 7A-C), in unadjusted or adjusted models. Beta diversity showed no significant differences between participants with minimal and elevated depressive symptoms (ANOSIM test, R = 0.0392, p = 0.210, Fig. 3**D-E**), nor between participants with minimal and elevated anxiety symptoms (ANOSIM test, R= −0.0157, p = 0.642, Supplementary Fig. 7D-E).

**Fig. 3.**
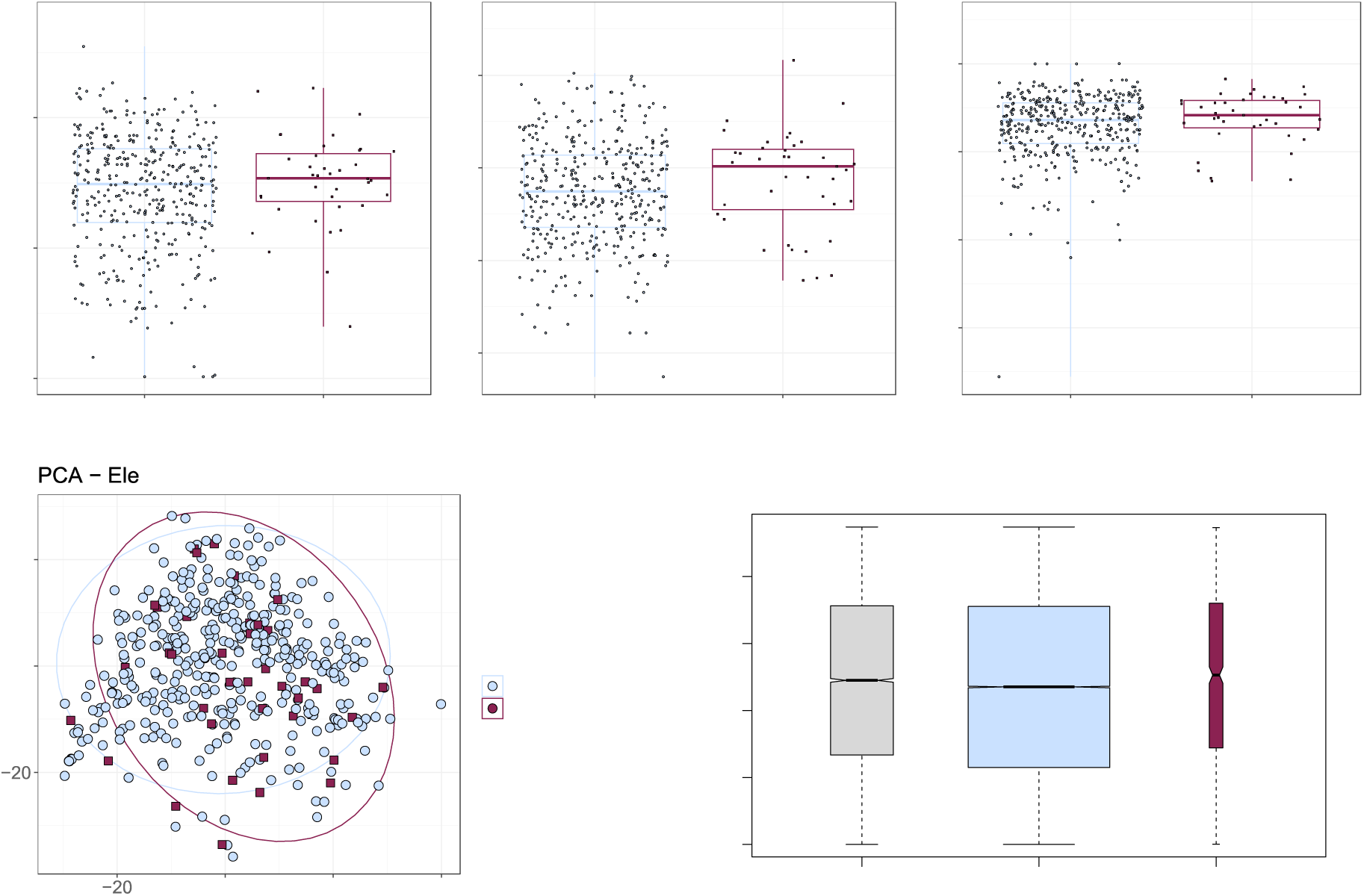
Alpha and beta diversities in participants with elevated and minimal depressive symptoms. (**A-C**) Quantification of the variation in richness (Chao index) and diversity (Shannon entropy and Simpson index) of the oral microbiota between participants with minimal versus elevated depressive symptoms. No evidence of difference in richness/diversity (α-diversity) between the two groups. (**D-E**) Principal component analysis (based on the Aitchison distance) between participants with minimal versus elevated depressive symptoms, and results of the related analysis of similarities (ANOSIM) comparing the oral microbiota between and within each group. No evidence of difference in dissimilarities (β-diversity) between the two groups.

### 3.6 Differential abundance analysis on all oral microbiota in the samples

#### Depressive symptoms

When controlling for pre-specified confounders (age, SES, diet, lifestyle risk, obesity, and hypertension), five oral species were positively associated with elevated depressive symptoms: *Streptococcus anginosus*, *Prevotella nigrescens ATCC 33563*, *Olsenella sp. oral taxon 807*, a *Campylobacter* uncultured bacterium, and a *Peptoniphilus* unidentified species (ORs > 1 with q < 0.1, Table 2). Notably, *S. anginosus* had an odds ratio of 3.39 with q < 0.05 (Table 2; Supplementary Fig. 8), approximately equivalent to Cohen’s d = 0.5 (i.e., a medium effect size) (Chen et al., 2010). Moreover, five oral species were inversely associated with elevated depressive symptoms: *Streptococcus cristatus*, *Eikenella sp. NML130454*, *Capnocytophaga gingivalis ATCC 3362*, *Corynebacterium sp. oral clone AK153* and *Streptococcus respiraculi* (ORs < 1 with q < 0.1, Table 2). The unadjusted differential abundance analysis for depression produced no significant result (all q > 0.1).

**Table 2.**
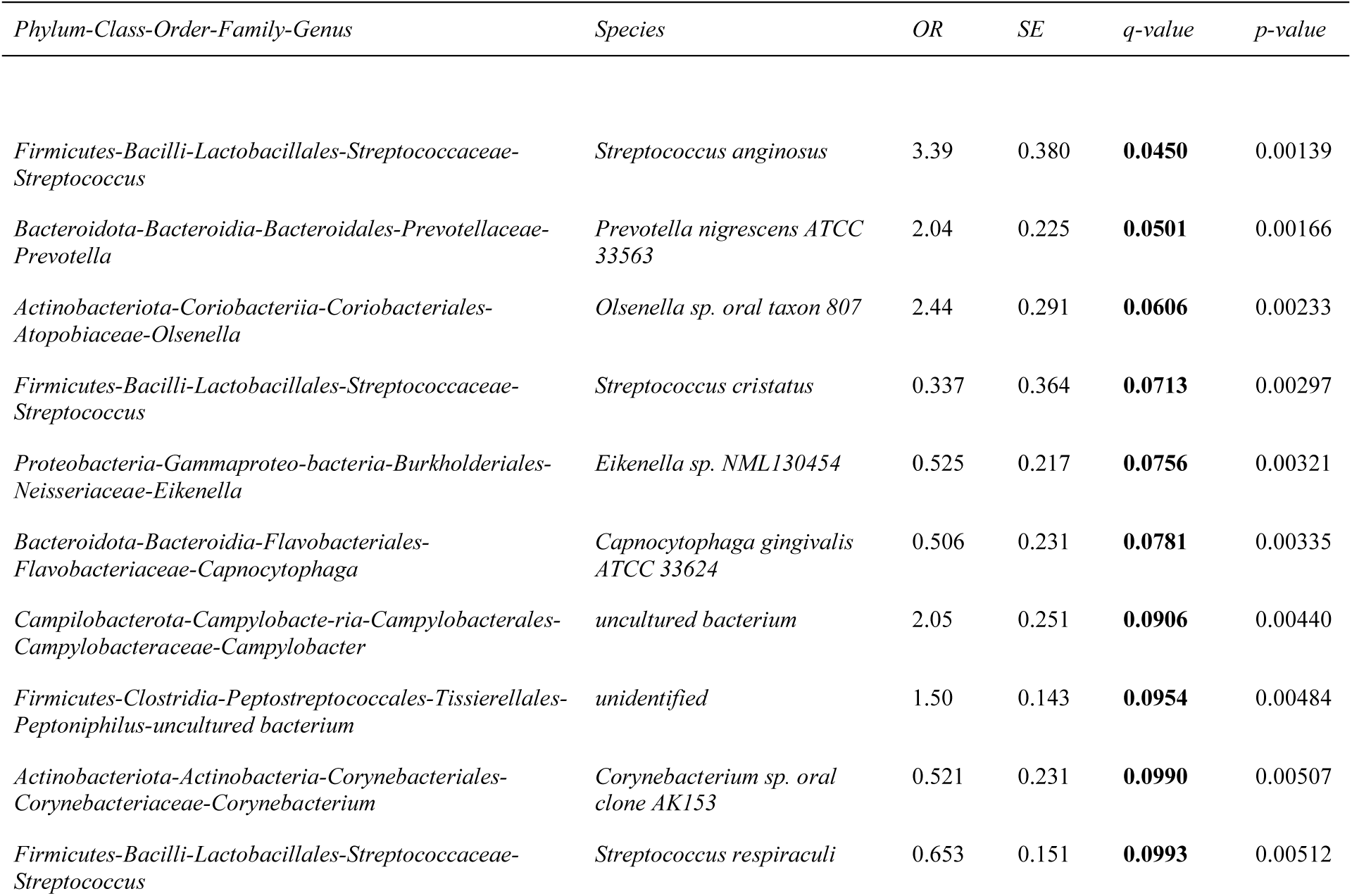
Significant results (q < 0.1) of the differential abundance analysis across all oral taxa at the species level for depressive symptoms, controlling for confounders. Species are ordered based on ascending q-values. All tests are adjusted for confounders: age, SES, diet, lifestyle risk, obesity, and hypertension. OR: odds ratio estimate; SE: standard error; q-value: the p-value after Benjamini-Hochberg adjustment for multiple hypothesis testing; SES, Socio-Economic Status.

#### Anxiety symptoms

Two species were differentially abundant between participants with minimal and elevated anxiety symptoms when controlling for age, SES, diet, lifestyle risk, obesity and hypertension: *Prevotella melaninogenica* and *Eikenella sp. NML130454* (q < 0.1, Table 3). These two oral species were inversely associated with elevated anxiety symptoms (both ORs < 1, Table 3).

**Table 3.** Significant results (q < 0.1) of the differential abundance analysis across all oral taxa at the species level for anxiety symptoms, controlling for confounders. Species are ordered based on ascending q-values. Both tests are adjusted for confounders: age, SES, diet, lifestyle risk, obesity, and hypertension. OR, Odds Ratio estimate; SE, Standard Error; q-value, the p-value after Benjamini-Hochberg adjustment for multiple hypothesis testing; SES, Socio-Economic Status.

| *Phylum-Class-Order-Family-Genus* | *Species* | *OR* | *SE* | *q-value* | *p-value* |
| --- | --- | --- | --- | --- | --- |
| *Bacteroidota-Bacteroidia-Bacteroidales-Prevotellaceae-Prevotella* | *Prevotella melaninogenica* | 0.641 | 0.145 | **0.0657** | 0.00242 |
| *Proteobacteria-Gammaprote-obacteria-Burkholderiales-Neisseriaceae-Eikenella* | *Eikenella sp. NML130454* | 0.602 | 0.175 | **0.0872** | 0.00399 |

## 4. Discussion

In this study of population-based men, we observed limited evidence for an association between our oral pathogen composite and elevated depressive, but not anxiety symptoms. We did not observe evidence of associations between oral microbiota alpha and beta diversity and depressive or anxiety symptoms. However, we did observe positive and inverse associations between various oral microbial taxa and both depressive symptoms and anxiety symptoms. Our analyses identified several oral species, *P. nigrescens, F. nucleatum subsp. animalis ATCC 51191, F. nucleatum subsp. animalis, F. nucleatum subsp. vincentii, S. anginosus, Olsenella sp. oral taxon 807*, a *Campylobacter* uncultured bacterium, and an unidentified species of an uncultured bacterium from the *Peptoniphilus* family, that were associated with elevated depressive symptoms. Conversely, several species showed an inverse association with depressive symptoms (*S. cristatus*, *Eikenella sp. NML130454*, *C. gingivalis ATCC 33624*, *Corynebacterium sp. oral clone AK15*3, and *S. respiraculi*), as well as anxiety symptoms (*P. melaninogenica* and *Eikenella sp. NML130454*).

There was a strong positive association between one of the oral species in our composite, *P. nigrescens* (*P. nigrescens ATCC 33563*) and elevated depressive symptoms. This finding is consistent with results from a recent study in patients with clinical depression, where *P. nigrescens* was more abundant in depressed participants (Wingfield et al., 2021). That study focused on young adults (18-38 years old), while our cohort comprised middle-aged and older participants (33-96 years old). Together, these results suggest that the association between *P. nigrescens* and elevated depressive symptoms may be relevant throughout adulthood. The pathogenicity of *P. nigrescens* includes its established link to periodontitis (Stingu et al., 2013) and previously reported immune responses *in-vivo* (Larsen, 2017). This association warrants further investigation in larger cohorts and longitudinal studies to better understand its potential causality.

Three subspecies of *F. nucleatum* were positively associated with elevated depressive symptoms (*subsp. vincentii*, *subsp. animalis ATCC 51191*, and *subsp. animalis*). Metagenomics enables the analysis of within-species differences, providing insights into potential differential effects of varying subspecies/strains (Escobar-Zepeda et al., 2015). Future research using metagenomic rather than 16S rRNA sequencing could further enhance our understanding of the complex relationships between *F. nucleatum* species and depressive symptoms and help determine whether these initial findings can be replicated.

Beyond the candidate oral pathogens, positive associations were found between several taxa and elevated depressive symptoms. These taxa were *S. anginosus*, *Olsenella sp. oral taxon 807*, a *Campylobacter* uncultured bacterium, and an unidentified species of an uncultured bacterium from the *Peptoniphilus* family. To our knowledge these are novel associations not previously reported by similar studies (Simpson et al., 2020; Wingfield et al., 2021). We note that *S. anginosus* bacteremia is associated with infections of the skin, soft tissue, and biliary tract (Su et al., 2018).

Interestingly, several oral species in in our study were inversely associated with elevated depressive symptoms: *S. cristatus*, *Eikenella sp. NML130454*, *C. gingivalis ATCC 33624*, *Corynebacterium sp. oral clone AK15*3, and *S. respiraculi*. Elsewhere, inverse associations between many oral taxa and clinical depression or depressive symptoms have been reported (Wingfield et al., 2021); however, none of the species previously identified overlapped with our results.

We report fewer associations between the oral microbiota and anxiety symptoms than observed for depressive symptoms. The species *P. melaninogenica* and *Eikenella sp. NML130454* showed an inverse association, where increased abundance was associated with lower anxiety symptoms. Depression and anxiety have different biological processes underpinning them. Inflammation is associated with an increased risk of clinical depression (Pasco et al., 2010) and is thought to contribute to the development and persistence of depression through mechanisms involving cytokine signaling, neurotransmitter metabolism, and hypothalamic–pituitary– adrenal axis dysregulation (Otte et al., 2016). Anxiety, on the other hand, is influenced by a more complex interplay of genetic, neurochemical, and structural factors (Craske et al., 2017), with inflammation playing a potential but less well-defined role to date.

We observed limited evidence for an association between a composite of oral bacteria, comprising *P. gingivalis* (Cunha et al., 2019; Dominy et al., 2019; Poole et al., 2013), *F. nucleatum* (Hajishengallis and Chavakis, 2021), *T. denticola* (Kin et al., 2020; Miklossy, 2011; Nieminen et al., 2018; Riviere et al., 2002; Simpson et al., 2020), and *P. nigrescens* (Wingfield et al., 2021), and elevated depressive symptoms. This method of composite score analysis accounts for the integrated effects of microbial functional guilds, rather than examining them in isolation (Kers and Saccenti, 2022; Quinn et al., 2019; Wu et al., 2021); this method is relatively recent, and none of the clinical studies on the oral microbiota in mental and neurodegenerative disorders discussed to date has employed this approach. Furthermore, there are no definitive guidelines for adapting a compositional data analysis approach to the development of a multiple-species composite score, and alternative methods for creating the composite could yield different results. Further refinement may be necessary to better understand which individual bacteria should be included in such a composite if this approach was to be pursued again.

We did not observe associations between either alpha or beta diversity and depressive or anxiety symptoms. This aligns with the findings in Australian young adults, where no disparities in alpha or beta diversity were reported between anxiety or depression groups (Simpson et al., 2020). However, other studies have presented contrasting results; for example, variations in alpha and beta diversity in the salivary microbiota were reported between individuals with higher psychological distress compared to controls (Kohn et al., 2020), and significant differences in beta diversity reported in the salivary microbiota of a clinically depressed cohort compared to controls (Wingfield et al., 2021).

This study represents a pioneering effort in understanding the association between oral microbiota and elevated common mental disorder symptoms. The study methods adopted a causal modelling framework, controlling for potential confounding variables drawn from the literature. The study sample was not selected on the basis of a disease, and the large sample size (436 participants) provided a robust control group. The high-quality of the oral microbiota data afforded information on 900 unique OTUs, some identified to the subspecies level. However, the study has several limitations. First, there were few “cases” of elevated depressive symptoms in our sample, which limited our statistical power and may have accounted for the weak evidence arising from our primary analysis. Depression and anxiety disorders are nearly twice as prevalent in women (Craske et al., 2017; Otte et al., 2016) and there may be disparities in oral microbiota between sexes (Lira-Junior et al., 2018). The male-only sample also limits generalizability of our findings to women. The use of the HADS questionnaire, while consistent with previous literature, does not provide clinical diagnoses of depressive or anxiety disorders.

Dichotomizing HADS scores for meaningful clinical representation may also have led to a general loss of statistical power in the study (increased type 2 error), while the cross-sectional, observational study design means that we cannot ascribe causality. Differences in oral bacteria across niches and study-specific swabbing protocols could introduce variability (Lamont et al., 2018). This limitation is not unique to this study; methodological heterogeneity is a feature of the oral and broader microbiota field at large (Mirzayi et al., 2021; Zaura et al., 2021). Regarding data collection, factors like mouthwash use, food or beverage consumption, and tooth brushing habits were not controlled, potentially influencing the oral microbiota composition (Minty et al., 2021; Sotozono et al., 2021). Finally, the study’s relatively narrow age range, with predominantly middle-aged and older individuals, may not reflect younger populations (Burcham et al., 2020; Willis et al., 2022), and the regional nature of the cohort may also limit generalizability (Sanders et al., 1999).

## Conclusion

This study provides new evidence showing associations between the oral microbiota and symptoms of common mental disorders, particularly for depressive symptoms. While several species were positively associated with elevated depressive symptoms, others had inverse associations with both elevated depressive and anxiety symptoms. Research into the pathogenic effect of oral pathogens has emerged and justified this study; however, more research replicating our findings and seeking clarity on the mechanistic pathways involved is needed, ideally in larger sample sizes of both sexes.

## Supplementary materials

Supplementary material associated with this preprint is included in the online version.

## Acknowledgements

We thank the participants of the Geelong Osteoporosis Study (GOS). The GOS was funded by the National Health and Medical Research Council (NHMRC) Australia (projects 299831, 251638, 628582).

We also acknowledge G. Giles of the Cancer Epidemiology Centre of The Cancer Council Victoria for permission to use the Dietary Questionnaire for Epidemiological Studies (Version 2), Melbourne: The Cancer Council Victoria, Australia, 1996.

## Author statement

**Data collection**: Julie A Pasco, Lana J Williams.

**Conceptualization/methodology**: Felice N Jacka, Amy Loughman, Samantha L Dawson, Fannie Kerff.

**Data analysis**: Fannie Kerff.

**Supervision**: Samantha L Dawson, Amy Loughman, Felice N Jacka.

**Writing – original draft**: Fannie Kerff, Samantha L Dawson, Amy Loughman.

**Writing – review & editing**: Fannie Kerff, Samantha L Dawson, Amy Loughman, Felice N Jacka, Julie A Pasco, Lana J Williams.

## Abbreviations

HADS: Hospital Anxiety and Depression Scale
CLR: Centered Log Ratio
OTUs: Operational Taxonomic Units
SES: Socio-Economic Status
IRSAD: Index of Relative Socio-economic Advantage and Disadvantage
E-DII: Energy-adjusted Dietary Inflammatory Index
OR: Odds Ratio
CI: confidence interval

