## Supplementary material for "Associations between oral microbiota pathogens and elevated depressive and anxiety symptoms in men"

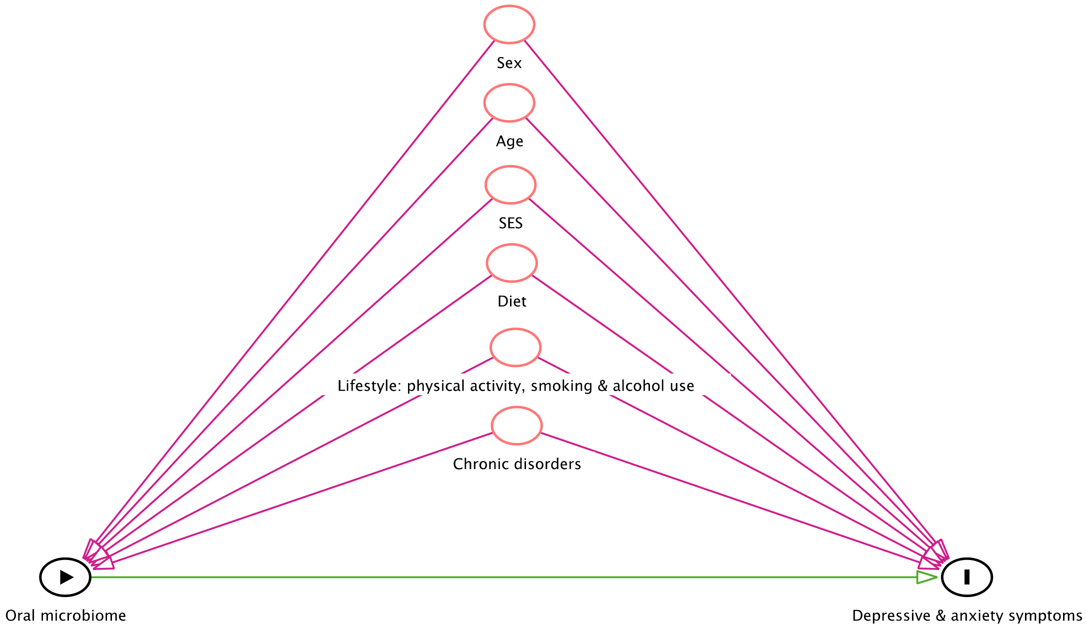


Supplementary Fig. 1. **Directed Acyclic Graph**.
*A priori* confounding variables in this analysis: sex, age, SES, diet, lifestyle risk (physical activity, smoking and alcohol use) and the presence of chronic disorders (especially, obesity and hypertension). SES, Socio-Economic Status.

| Component variable | No risk (0) | Low risk (1) | High risk (2) |
|---|---|---|---|
| Smoking | Never | Former | Current |
| Current alcohol intake | Abstinence | Modest | Heavy |
| Physical activity | Active | Inactive | |

*Notes*. Definition of the LRS component variables: smoking, current alcohol intake and physical activity.

Supplementary Table 1. **Measure of the lifestyle risk score**.
The score ranges from 0 to 5; a score of 5 corresponds to the maximum risk. This combines information on smoking status, current alcohol intake and exercise in the participants. Smoking status in the individuals was defined as never (0; no risk), former (1; low risk), or current (2; high risk). Current alcohol intake included three groups: abstinent i.e., no drinks/week (0; no risk), modest i.e., 1–7 drinks/week (1; low risk), and heavy i.e., > 7 drinks/week (2; high risk) drinkers. A drink was defined as 14 grams of alcohol (National Institute of Alcohol Abuse and Alcoholism). Finally, physical activity was defined as active (0; no risk) or inactive (1; low risk). Physical activity was computed using a combination of the Baecke (< 60 years-old (Baecke et al., 1982)) and Voorrips (>60 years-old (elderly) (Voorrips et al., 1991)) questionnaires dichotomized at the median.

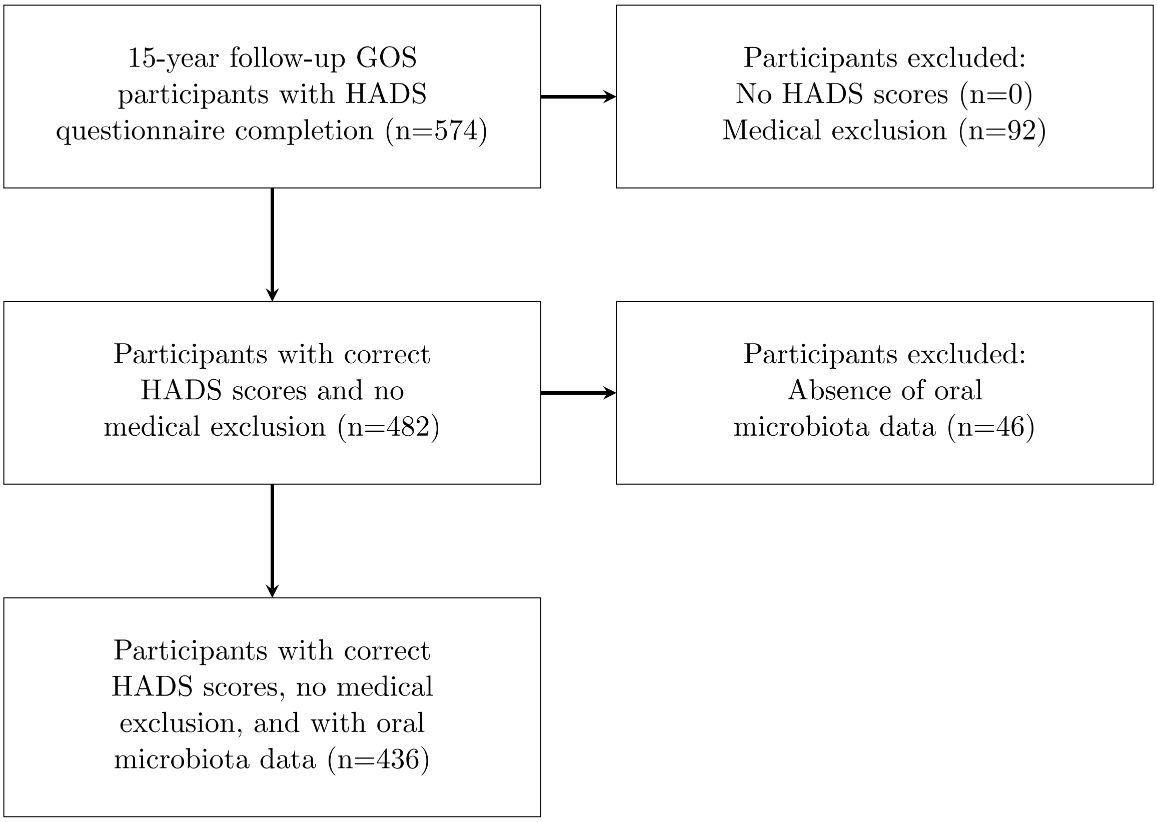


Supplementary Fig. 2. **Flow chart of the inclusion and exclusion process of the participants**. Description of the selection of the 436 participants included in the analysis. GOS, Geelong Osteoporosis Study; HADS, Hospital Anxiety and Depression Scale.

| | Past-year conditions | | | | | Long-term conditions | | *Total (omitting duplicates)* |
|---|---|---|---|---|---|---|---|---|
| | Cancer | Severe heart disease | Severe brain disease | Severe liver disease | Severe kidney disease | Brain conditions | Heart conditions | |
| | Lung, bowel, breast, uterus, cervix, throat, brain, melanoma, non-melanoma skin, leukemia, myeloma, lymphoma, prostate, bone marrow lymphoma, clear cell carcinoma of the kidney, pituitary gland tumor | Heart attack (myocardial infarction), stroke, heart disease, heart bypass surgery, heart pacemaker insertion, stents in heart, leaky or defects in valve and aorta, valve replacement, myocardial bridging and ectopic heartbeat, hole in the heart | Brain damage, brain injury, brain aneurysm, and subarachnoid hemorrhage | Liver hepatic failure and cirrhosis liver | Kidney renal failure | Parkinson's disease, Alzheimer's disease | Hole in heart since birth | |
| *Total* | 53 | 31 | 4 | 2 | 4 | 3 | 2 | 92 (16%) |

Supplementary Table 2. **Severe medical conditions excluded from the analysis**.

List of the medical conditions that led to the exclusion of the participants. 92 participants (16%) were excluded in total.

| Species | Prevalence (# of samples) | Prevalence (% of samples) | Correlation with the composite | Included in pathogen composite? |
|---|---|---|---|---|
| *Fusobacterium nucleatum subsp. polymorphum* | 390 | 89.4 % | 0.5 | Yes |
| *Fusobacterium nucleatum subsp. vincentii ATCC 49256* | 372 | 85.3 % | 0.8 | Yes |
| *Fusobacterium nucleatum subsp. animalis* | 364 | 83.5 % | 0.8 | Yes |
| *Fusobacterium nucleatum subsp. vincentii* | 327 | 75.0 % | 0.7 | Yes |
| *Fusobacterium nucleatum subsp. animalis ATCC 51191* | 287 | 65.8 % | 0.7 | Yes |
| *Treponema denticola* | 193 | 44.3 % | 0.5 | Yes |
| *Fusobacterium nucleatum subsp. animalis 7_1* | 177 | 40.6 % | 0.5 | Yes |
| *Prevotella nigrescens ATCC 33563* | 171 | 39.2 % | 0.4 | Yes |
| *Fusobacterium nucleatum subsp. vincentii 3_1_36A2* | 71 | 16.3 % | 0.2 | Yes |
| *Fusobacterium nucleatum subsp. fusiforme ATCC 51190* | 51 | 11.7 % | *0* | *No* |
| *Porphyromonas gingivalis* | 47 | 10.8 % | 0.1 | Yes |
| *Aggregatibacter actinomycetemcomitans D7S-1* | 10 | *2.29 %* | *-0.2* | *No* |
| *Aggregatibacter actinomycetemcomitans DSM 8324* | 10 | *2.29 %* | *-0.3* | *No* |
| *Aggregatibacter actinomycetemcomitans* | 2 | *0.46 %* | *-0.4* | *No* |
| *Klebsiella pneumoniae* | *0* | *0 %* | */* | *No* |

Supplementary Table 3. **Candidate oral species for the pathogen composite: prevalence in the samples and correlation with the composite.**

Prevalence reported both as raw number of samples and as percentage of samples the species are detected in. Correlations between each species and the pathogen composite are reported.

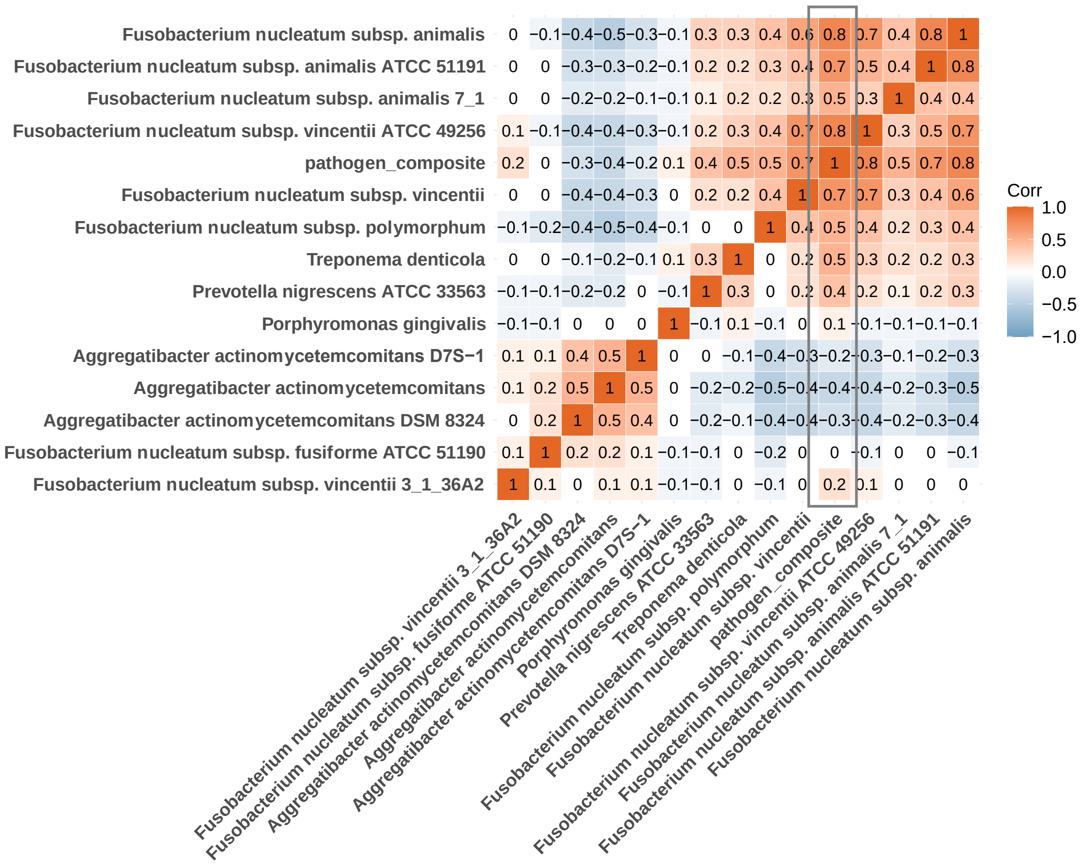


Supplementary Fig. 3. **Correlation matrix of the species within the pathogen composite.**
The numbers represent the correlation coefficients of species CLR-transformed abundances. All three species of *Aggregatibacter actinomycetemcomitans* and species *Fusobacterium nucleatum subsp. fusiforme ATCC 51190* are not positively correlated with the pathogen composite (correlation ≤ 0). CLR, Centered Log Ratio.

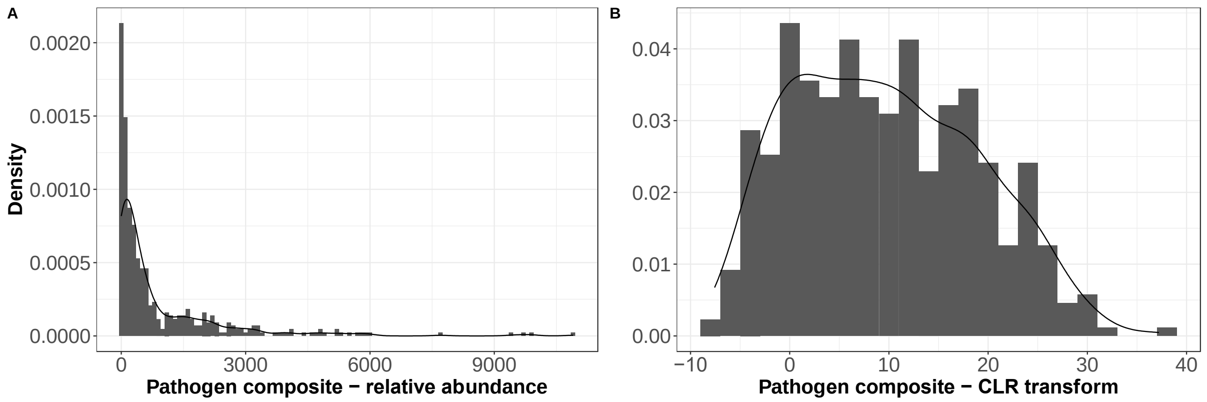


Supplementary Fig. 4. **Histograms and density curves of the distributions of the oral pathogen composite.**

**A** Relative abundance. **B** CLR transformed abundance. The pathogen composite comprises species *P. gingivalis*, *F. nucleatum* (excluding *F. nucleatum subsp. fusiforme ATCC 51190*), *T. denticola*, and *P. nigrescens*. CLR, Centered Log Ratio.

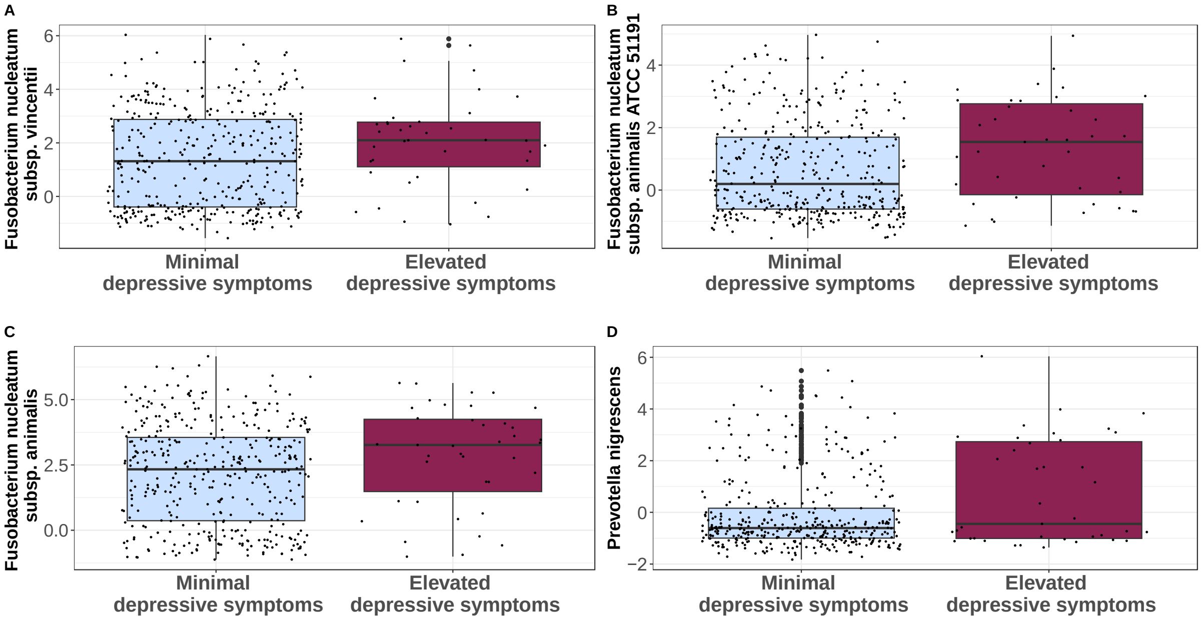


Supplementary Fig. 5. **Distributions of species in participants with minimal and elevated depressive symptoms (CLR transformed abundance).**

**A** *F. nucleatum subsp. vincentii*. **B** *F. nucleatum subsp. animalis ATCC 51191*. **C** *F. nucleatum subsp. animalis*. **D** *P. nigrescens.* The different distributions in the sub-samples illustrates the significantly greater abundances of *F. nucleatum subsp. vincentii*, *F. nucleatum subsp. animalis ATCC 51191*, *F. nucleatum subsp. animalis*, and *P. nigrescens* species in participants with elevated depressive symptoms. CLR, Centered Log Ratio.

| | | OR | SE | CI (95%) | p-value |
|---|---|---|---|---|---|
| Unadjusted models | | | | | |
| A1 | *P. gingivalis* | 0.821 | 0.167 | 0.564 - 1.10 | 0.237 |
| A2 | *T. denticola* | 1.16 | 0.127 | 0.894 - 1.47 | 0.255 |
| A3 | *F. nucleatum subsp. vincentii ATCC 49256* | 1.08 | 0.135 | 0.830 - 1.41 | 0.575 |
| A4 | *F. nucleatum subsp. vincentii 3_1_36A2* | 0.799 | 0.163 | 0.560 - 1.07 | 0.170 |
| A5 | *F. nucleatum subsp. vincentii* | 1.21 | 0.133 | 0.932 - 1.57 | 0.152 |
| A6 | *F. nucleatum subsp. animalis 7_1* | 0.885 | 0.142 | 0.659 - 1.16 | 0.392 |
| A8 | *F. nucleatum subsp. animalis ATCC 51191* | 1.27 | 0.128 | 0.983 - 1.63 | 0.0645 |
| A9 | *F. nucleatum subsp. animalis* | 1.21 | 0.135 | 0.930 - 1.58 | 0.157 |
| A7 | *F. nucleatum subsp. polymorphum* | 1.06 | 0.135 | 0.817 - 1.39 | 0.657 |
| A10 | *P. nigrescens ATCC 33563* | 1.19 | 0.124 | 0.922 - 1.50 | 0.168 |
| Adjusted models | | | | | |
| B1 | *P. gingivalis* | 0.813 | 0.166 | 0.559 - 1.09 | 0.215 |
| B2 | *T. denticola* | 1.14 | 0.130 | 0.880 - 1.47 | 0.298 |
| B3 | *F. nucleatum subsp. vincentii ATCC 49256* | 1.05 | 0.141 | 0.798 - 1.39 | 0.730 |
| B4 | *F. nucleatum subsp. vincentii 3_1_36A2* | 0.816 | 0.167 | 0.567 - 1.10 | 0.225 |
| B5 | *F. nucleatum subsp. vincentii* | 1.19 | 0.137 | 0.909 - 1.56 | 0.204 |
| B6 | *F. nucleatum subsp. animalis 7_1* | 0.867 | 0.145 | 0.642 - 1.14 | 0.325 |
| B8 | *F. nucleatum subsp. animalis ATCC 51191* | 1.28 | 0.134 | 0.981 - 1.66 | 0.0665 |
| B9 | *F. nucleatum subsp. animalis* | 1.18 | 0.141 | 1.894 - 1.56 | 0.246 |
| B7 | *F. nucleatum subsp. polymorphum* | 1.06 | 0.140 | 0.809 - 1.40 | 0.667 |
| B10 | *P. nigrescens* | 1.19 | 0.129 | 0.914 - 1.52 | 0.186 |

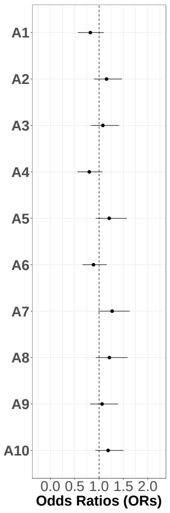


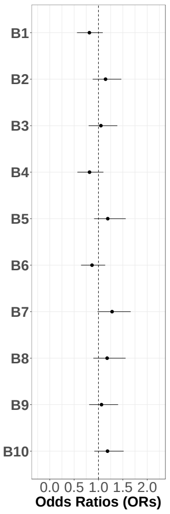


Supplementary Fig. 6. **Logistic regressions result for species within the composite and elevated anxiety symptoms.**

No significant association with any species. Adjusted models control for confounders: age, SES, diet, lifestyle risk, obesity and hypertension. OR, Odds Ratio estimate; SE: Standard Error; CI (95%): 95% Confidence Interval; SES, Socio-Economic Status.

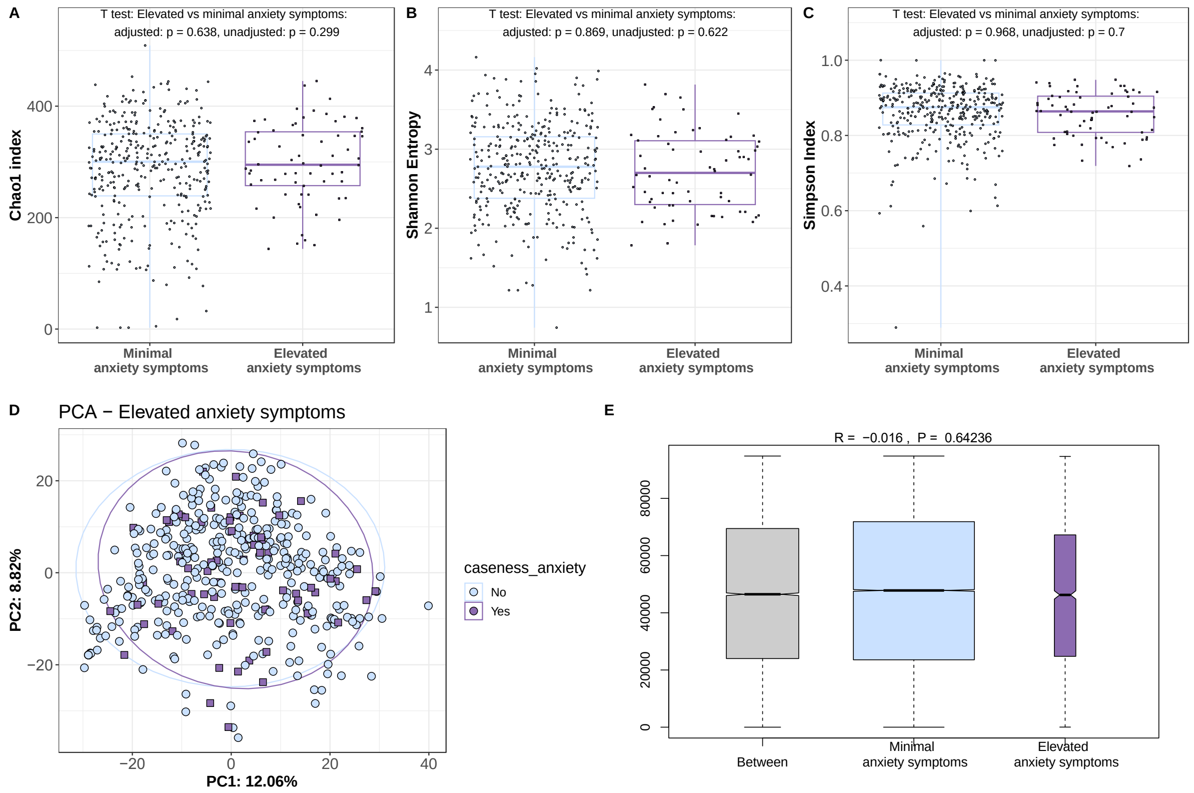


Supplementary Fig. 7. **Alpha and beta diversities in participants with elevated and minimal anxiety symptoms.**

(**A-C**) Quantification of the variation in richness (Chao index) and diversity (Shannon entropy and Simpson index) of the oral microbiota between participants with minimal versus elevated anxiety symptoms. No evidence of difference in richness/diversity (α-diversity) between the two groups. (**D-E**) Principal component analysis (based on the Aitchison distance) between participants with minimal versus elevated anxiety symptoms, and results of the related analysis of similarities (ANOSIM) comparing the oral microbiota between and within each group. No evidence of difference in dissimilarities (β-diversity) between the two groups.

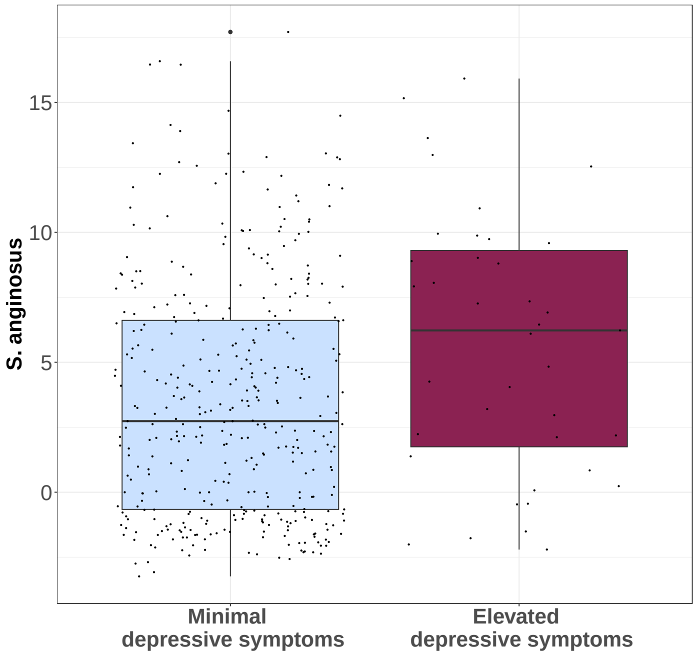


Supplementary Fig. 8. **Distributions of *Streptococcus anginosus* in participants with minimal and elevated depressive symptoms (CLR transformed abundance).**

The large difference in the medians of both samples illustrates this significantly greater abundance of *S. anginosus* in the participants with elevated depressive symptoms. CLR, Centered Log Ratio.
